# Thermoconforming rays of the star-nosed mole

**DOI:** 10.1101/2022.11.18.517115

**Authors:** Glenn J. Tattersall, Kevin L. Campbell

## Abstract

The star-nosed mole (*Condylura cristata*) is well known for its unique star-like rostrum (‘star’) which is formed by 22 nasal appendages highly specialised for tactile sensation. As a northerly distributed insectivorous mammal occupying both aquatic and terrestrial habitats, this sensory appendage is regularly exposed to cold water and thermally conductive soil, leading us to ask whether the surface temperature, a proxy for blood flow to the star, conforms to the local ambient temperature to conserve body heat. Alternatively, given the high functioning and sensory nature of the star, we posited it was possible that the rays may be kept continually warm when foraging, with augmented peripheral blood flow serving the metabolic needs of this tactile sensory organ. To test these ideas, we remotely monitored the surface temperatures of the star and other uninsulated appendages in response to changes in local water or ground temperature in captive, wild-caught star-nosed moles. While the tail responded to increasing heat load through vasodilation, the surface temperature of the star consistently thermoconformed, varying passively in surface temperature, suggesting little evidence for thermoregulatory vasomotion. This thermoconforming response may have evolved as a compensatory response related to the high costs of heat dissipation to water or soil in this actively foraging insectivore.

**Summary Statement (for JEB Submission):** The highly mechanosensitive nasal rays of the star-nosed mole conform closely with ambient temperature thereby minimizing heat loss without apparent changes in sensory performance.

## Introduction

The star-nosed mole (*Condylura cristata*) is a highly specialized insectivore that is uniquely adapted to foraging in both terrestrial and aquatic habitats (Catania, 1999; Catania, 2000). Found throughout eastern North America, and extending northward to the southern limit of permafrost, this nearly blind predator primarily relies on its incredibly touch sensitive nasal appendages to rapidly identify and consume hundreds of tiny prey items per day to fuel its high rate of metabolism (Campbell et al., 1999; Catania and Remple, 2005). Owing to its distinctive morphology, the eponymous nose (‘star’) of the star-nosed mole has been extensively studied for its sensory functions (Catania, 2000; Gould et al., 1993; Sachdev and Catania, 2002; Sawyer et al., 2014). The rostrum houses 22 separate rays, 11 per side, that sample the tactile environment 10-15 times per second while foraging (Gerhold et al., 2013). The star is highly vascularized with two large non-muscularised blood sinuses occupying approximately 40% of the volume of each ray. Capillaries are also evident throughout the dermis underlying the thousands of sensory papillae (Eimer’s organs) within each ray (van Vleck, 1965).

The star acts as the primary mechanosensory organ, with >100,000 myelinated nerve fibres innervating the roughly 30 thousand Eimer’s organs covering the surface of the nose (Catania, 1995; Catania, 1999). While all rays contribute to the remarkable tactile acuity of the star, the inner most 11^th^ ray serves as a mechanosensory fovea (Catania, 2011; Sachdev and Catania, 2002), and when foraging, moles will redirect their attention to allow this appendage to investigate stimuli immediately prior to consumption. The speed of this behaviour is astonishing, as star-nosed moles can locate, identify, and ingest prey items in as little as 102 ms, crowning them as one of fastest eaters of the animal kingdom (Catania and Remple, 2005). In principle, the sensory structures of endotherms are metabolically active and highly temperature sensitive tissues that are expected to function more effectively when maintained at a warm and stable temperature (Glaser and Kroger, 2017). The elephant’s (*Loxodonta africana*) trunk, for example, is the warmest part of the skin (Weissenboeck et al., 2010). During development, sensory nerves provide a map for arterial growth through secretion of VEGF (Mukouyama et al., 2002), and thus there may be a natural tendency for sensory activity to be associated with changes in blood flow, leading to potential trade-offs between sensory response and energy supply (in the form of body heat). Indeed, a precedence for thermal-sensory associations exists (Glaser and Kroger, 2017). For example, the sensory vibrissae of seals have been shown to maintain high temperatures owing to the high vascularity serving the underlying metabolically active sensory tissue and do not demonstrate vasoconstriction in the cold (Dehnhardt et al., 1998). This continuously elevated temperature helps maintain tactile sensitivity across a range of water temperatures that might otherwise hamper neuron function. Similarly, the eye heater organ found in billfish (Carey, 1982) has been argued to have evolved as a means of enhancing central nervous system functionality and enhanced sensory acuity in cold environments (Fritsches et al., 2005). The enhanced visual acuity gained from maintaining the eye at an elevated temperature provides a distinct reaction time advantage over their highly active, but ectothermic prey. On the other hand, consider an ectothermic predator that senses heat; the pit organ of the pit viper is a highly evolved infrared sensing tissue, consisting of a thin membranous tissue dense in mitochondria, myelinated and unmyelinated nerves, and a high vascularity (Goris et al., 2007; Hisajima et al., 2002). The latter has been argued to aid in providing rapid blood flow to help reduce after images formed during heat sensing (Goris et al., 2007). Intriguingly, the infrared sensing function of this tissue appears to operate better at cooler temperatures (Cadena et al., 2013), although this may be related to the nature of the signal transfer rather than the sensory tissue itself. Combined, there is reasonable precedence to expect that the nasal epidermal tissue of *C. cristata* would show elevated temperatures based on high rates of blood flow supporting the underlying metabolically active nervous tissues.

Endotherms typically have non-insulated or poorly insulated peripheral appendages that tend to exhibit strong vasomotor control, reflective of their involvement in redistributing body heat from the core to the periphery to aid in heat loss or to retaining heat in the core to aid in heat conservation (Erdsack et al., 2012; Tattersall et al., 2012; Weissenboeck et al., 2010). Thermally, these changes in blood flow can be assessed by examining how these uninsulated appendages’ surface temperatures change under different heat loads. This methodology has revealed classic examples of adjustable thermal radiators that include elephant ears (Phillips and Heath, 1992), the toucan bill (Tattersall et al., 2009), and rodent tails (Rand et al., 1965), to name a few. These surfaces tend to contribute greatly to the body’s capacity to dissipate heat, but are also subject to emotional influences, such as fear-induced vasoconstriction (Herborn et al., 2015; Vianna and Carrive, 2005). In subterranean species, conductive heat loss to the soil is particularly high from structures in direct contact with the substrate (see Plestilova et al., 2020). Given that the shallow surface tunnels of star-nosed moles are typically excavated in water saturated soils, heat transfer from the naked sensory appendages of the mole’s star is potentially extensive. This is especially germane during the winter months when aquatic foraging by this species is more prevalent (Campbell et al., 1999; Hamilton, 1931). Accordingly, due to the amphibious life history of the star-nosed mole and the highly specialised sensory nature of its nasal rays, sustaining high rates of warm arterial blood flow to the star may cause rapid heat loss and be energetically expensive for the star-nosed mole to maintain.

We thus tested whether star-nosed moles keep their nasal sensory appendages warm through vasodilation when exposed to the cold (thereby incurring high energetic costs) versus an energy conservation hypothesis wherein despite its high vascularity, the uninsulated star will show mainly passive warming and cooling responses (*i*.*e*., thermoconformation). We did this by exposing moles to different water temperatures and examining the patterns of surface temperatures from their potential thermal windows (eye, tail, limbs, and nasal rays).

## Materials and Methods

### Animal Handling

Three juvenile star-nosed moles of unknown sex were captured using Sherman and pitfall traps in the Willard Lake region, Ontario (49° 49’ 41.9088’’ N, 93° 58’ 5.2896’’ W) in June 2022 under the authorization of an Ontario Ministry of Natural Resources Wildlife Scientific Collector’s permit (1101339). During their time in captivity (21 days), each mole was housed within a two-chambered Rubbermaid system; one 76-L chamber, filled with moist soil to a depth of ~15 cm housed a small wooden nesting enclosure that was connected via plastic (ABS) piping to a second 76-L feeding chamber containing water to ~0.5 cm depth. Moles were allowed to access to soil via an ABS tee wye attached to the nesting enclosure. Moles were fed commercially sourced night crawlers (14-16 per day per animal) supplemented by wild-caught earthworms and other invertebrates from the site of collection. The nesting chambers were cleaned/replaced daily, while the feeding containers were thoroughly washed 2-3 times daily. All procedures were approved by the Ontario Ministry of Northern Development, Mines, Natural Resources and Forestry Wildlife Animal Care Committee (Protocol #22-493).

### Thermal Manipulations

Pilot observations conducted in air suggested that the nasal rays of the star-nosed mole were heterothermic and closely corresponded to surface/air temperatures. We thus experimentally exposed moles to warm (~30-32°C) or cold (2-4°C) water to elicit maximal thermal responses. Briefly, individual moles were transferred to a clean 19-L Rubbermaid™ container held at room temperature (17-20°C) for 10 minutes (post-handling period), exposed to warm or cold water at a depth of 1.5 cm for 10 minutes (water foraging period), and subsequently transferred to a dry container for a further 10-minute recovery period in air at room temperature. Each mole underwent this 30-minute procedure twice, once for each of the warm or cold-water challenges. During the 10-min water exposure, moles were periodically provided with earthworms (which were rapidly located and consumed regardless of water temperature) to encourage natural underwater foraging behaviours.

### Thermal Imaging

Time lapse infrared thermal imaging videos were captured every second with FLIR Research Studio software using a FLIR A8581 Mid-Wave Infrared Camera (resolution 1280×1024, 25 mm lens, thermal sensitivity <25 mK, accuracy ±1°C). The camera was mounted ~0.5 m above the animal providing a full view throughout the measurement period. We assumed emissivity of 0.95 and set object parameter settings in the software to the local air temperature (~17-20°C). From the captured videos, we extracted still frames at various time points throughout the post-handling, water foraging, and recovery periods. These still frames were extracted as 32-bit TIF files and exported for analysis in FIJI/ImageJ (Schindelin et al., 2012). Regions of interest were drawn over the front limbs, nose, nasal rays, eye, and tail using the free-hand tool, and the average temperature for each region was extracted from each still frame. Sample thermal images are depicted in Fig. 1.

**Figure 1.**
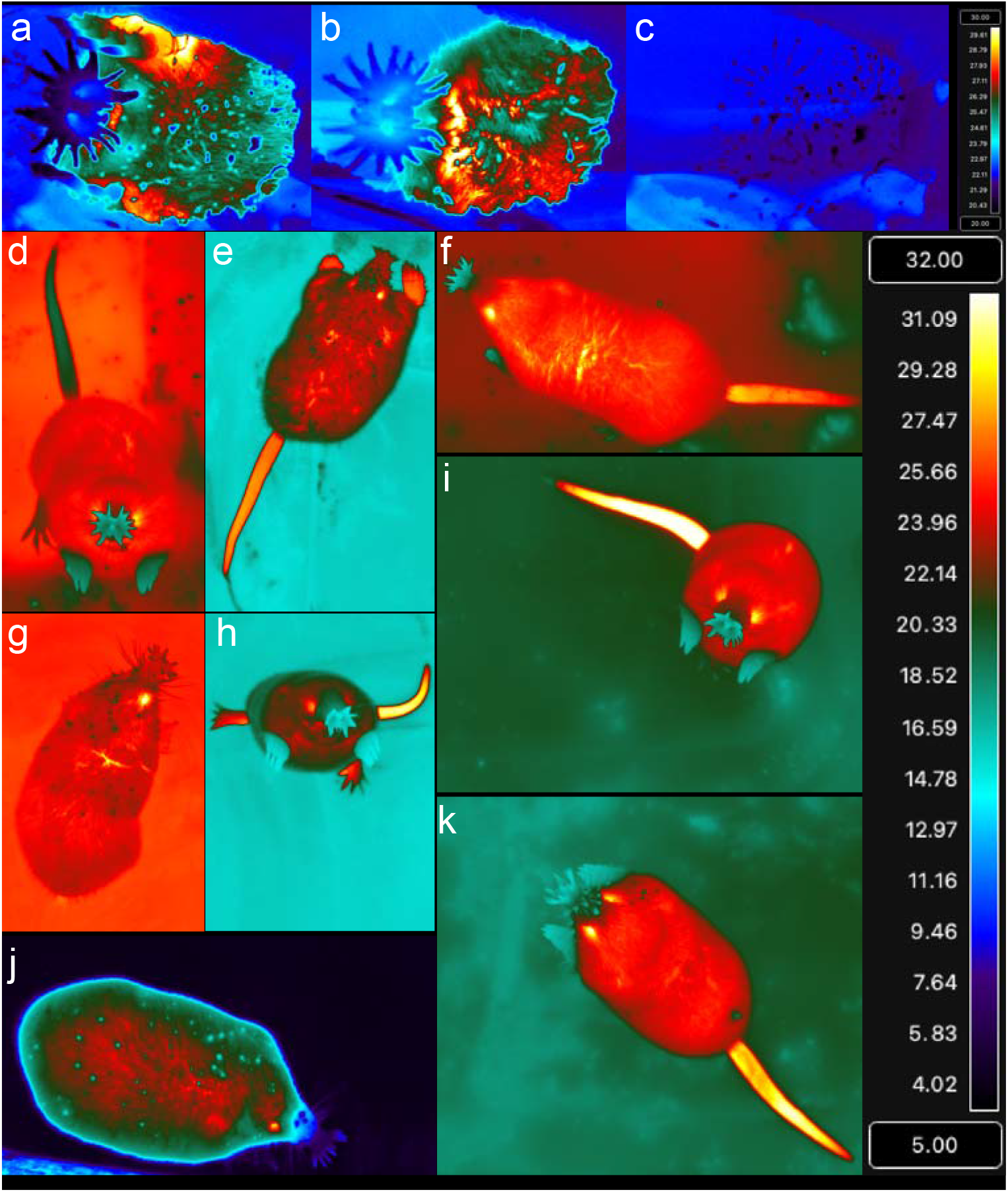
Representative thermal images of the star-nosed mole. Close-up image through a thin polyethylene sheet showing the warmer nostrils and distinctly cooler nasal rays when splayed out in surface contact (a,b) and the cool impressions (c) left behind at room temperature. Unless otherwise noted, the remaining images were captured during/following exposures to warm (~28-30°C) and cold (~2-4°C) environments. In d), immediately following handling after being placed into warm environment, e) immediate recovery at room temperature after exposed to warm water showing warm limbs, f) prior to being exposed to cold water while at room temperature, g) while foraging in warm water, h) recovery from exposure to warm water, i) and k) exposure to room temperature ground conditions, and j) during foraging in cold water. Note how the nasal ray and forelimb temperatures conform to ambient temperature while the eye and tail surface temperatures are generally much warmer. Temperature scale in upper right pertains to images a-c, while the larger temperature scale on the right pertains to images d-k.

### Data Analysis

Acknowledging that the small sample size limits broad conjectures based on biologically distinct replicates, we endeavoured to draw inferences of how various body part surface temperatures differed from prevailing ground or water temperatures based on biophysical principles outlined in Tattersall (2016). Surfaces that receive little blood flow or are insulated from warm blood (*i*.*e*., fur) are expected to be similar to local ground or water temperatures, and due to principles of thermoconformity, should have slopes close to 1. This hypothesis was tested using simple pairwise t tests (*P* values corrected using Bonferroni procedures for multiple hypotheses). Surfaces that have high and non-varying blood flow are expected to deviate from local temperature and to have a low slope (<<1) with respect to local temperature (tested by model comparison to a model where the slope is set to 1 using the offset function in R). Vasoactive body surfaces (*i*.*e*., adjustable thermal windows), would differ from local temperature when warm and exhibit a non-linear relationship with respect to local temperature, especially if vasoconstricted in the cold and vasodilated under warm conditions. While the slope between surface and local temperature may be less than 1, the obvious departure from linearity (tested by comparing whether the more complex model significantly reduces the residual sums of squares via a likelihood ratio test) reflects the vasoactive nature of the body surface. Statistical analyses were conducted in R (version 4.2.0).

## Results

Body surface temperatures of star-nosed moles were dependent on the ground and water temperature but in varying manners (Fig. 1). The nasal rays and front limbs were primarily thermoconforming body surfaces (Fig. 2 and Fig. 3) while the surface temperatures of the tail and eyes tended to be elevated. Eye surface temperatures differed significantly from local temperature (*t*_47_=9.4; *P*=9.03×10^−12^) and exhibited a slope significantly lower than 1 (χ^2^_df=1_ = 148; *P*<2×10^−16^). The star and front limb temperatures were not significantly different than local ambient temperatures (*t*_47_=-1.76; *P*=0.39 and *t*_47_=-8.55×10^−5^; *P*=1, respectively), showing a mostly linear, thermoconforming relationship. The tail was the most variable surface, being significantly warmer than local temperature (*t*_47_=4.25; *P*= 0.000580) but also exhibiting a non-linear relationship with local ground and water temperatures (χ^2^_df=1_ = 8.9; P=0.0029). Indeed, this surface was warmest at mid-range (~20°C) temperatures and thermoconforming at higher and lower temperatures (Fig. 3). While searching for food at room temperature, the nasal rays can be seen to be close to the temperature of the ground surface (Movie 1 and 2). We only once observed what might be evidence of vasodilation in the star that occurred during a prolonged, relaxed grooming session (Movie 3).

**Figure 2.**
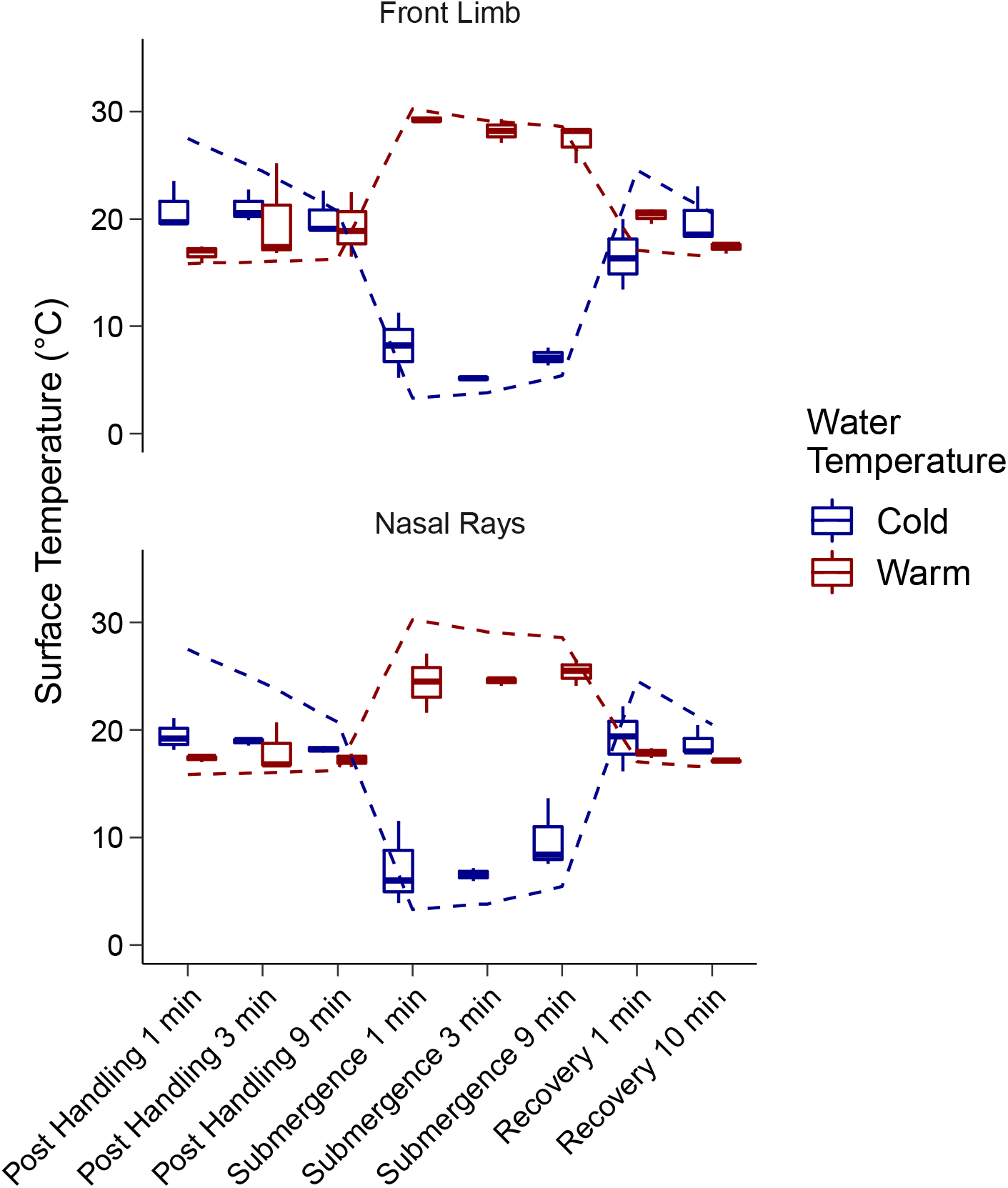
Surface temperatures of the front limbs and nasal rays (foraging appendages) prior, during, and following foraging in either cold (2-4°C) or warm (28-30°C) water. Symbols represent box and whisker plots (N=3), while the dotted lines represent the mean ground or water temperature for the respective cold or warm water exposures.

**Figure 3.**
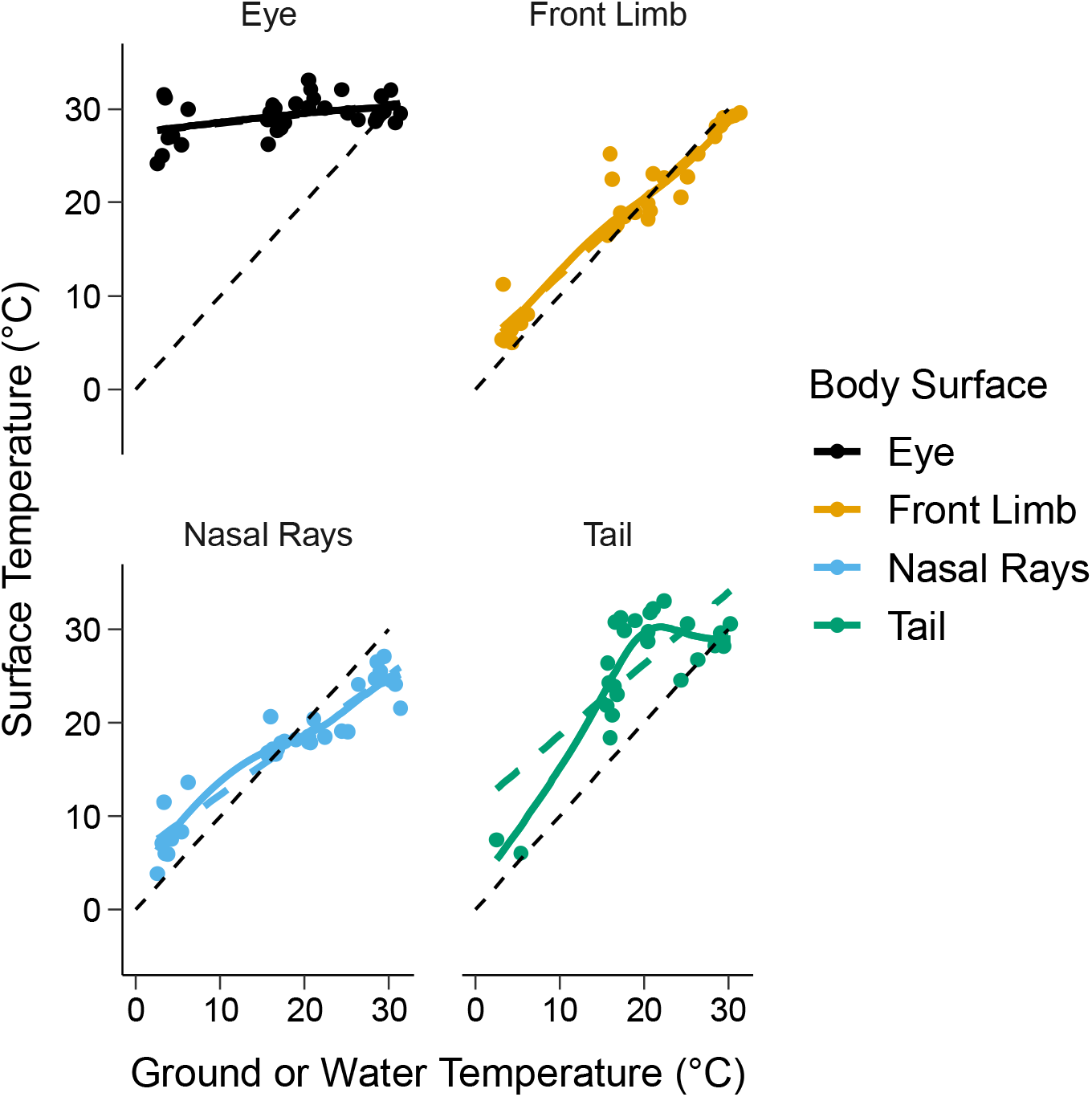
Surface temperature relationships of exposed star-nosed mole surfaces relative to ground or water temperatures across all measurement intervals. The front limbs and nasal rays are shown to mainly be thermoconforming surfaces. Eye surface temperature remained warm across all temperatures while the tail showed a complex non-linear relationship reflective of vasoconstriction in the cold and vasodilation at higher temperatures. Curves are included to illustrate the non-linearity of most surface temperatures. Bold dashed lines depict the linear regressions through the data, while the black dotted lines depict the line of equality.

## Discussion

We demonstrated using surface temperature measurements that the nasal rays and forelimbs of the star-nosed mole largely thermoconform to local water and ground temperatures, providing support for an energy conservation role for the star, whereas the tail acts as an adjustable thermal window, typical of many small mammals (Meyer et al., 2017; Rand et al., 1965). For the most part then, the star acts passively in terms of vasomotor responsiveness, with little evidence of vasodilation under heating scenarios; indeed, the nasal rays appear to be even a little cooler than expected at the higher ambient temperatures, a response well characterised in vampire bats (Kürten and Schmidt, 1982), canids (Balint et al., 2020), and numerous carnivores (Glaser and Kroger, 2017).

Owing to linkages between skin temperature and tactile sensitivity, it has been argued that cold rhinaria are incompatible with a mechanosensory role in mammals (Glaser and Kroger, 2017). Indeed, it is not unusual to expect warmer sensory structures to function more efficiently given what has been described in facultatively endothermic animals (Carey, 1982; Fritsches et al., 2005). However, canine olfactory-based tracking behaviours are enhanced at lower temperatures and higher humidity’s (Jinn et al., 2020), although this response has not been linked to their typically cool nose temperature. Thus, the strikingly lower temperature of the star relative to body temperature (37.7°C; Campbell et al., 1999) begs the question of whether this trait aids hinders the somatosensory function of the nasal rays while foraging. While not systematically studied, observed reductions in star surface temperatures did not appear to be associated with attendant reductions in prey detection ability as moles were able to rapidly locate and consume added prey items in both warm and cold water. The bigger question then is how does the star maintain high sensitivity/acuity in the cold if elevated blood flow and/or temperatures are not involved? An important caveat here is that epithelial cells are the dominant tissue of the rays, with much of the remaining volume composed of blood vessels and large sinuses (van Vleck, 1965). Although thousands of nerve fibres are interspersed within each ray, the cell bodies are located centrally within dorsal root ganglia. Accordingly, an argument for high metabolic requirements demanding high blood flow (*e*.*g*., Dehnhardt et al., 1998) seems inadequate in the case of the star-nosed mole. However, hints regarding the thermoconforming nature of the nasal star may be found in the specialised evolution of the sensory nerves innervating the nose. The mole’s nasal sensory nerves are of trigeminal ganglion origin and enriched in the expression of ion channels involved in innocuous mechanosensation (CNGA2, CNGA3, CNGA4, and FAM38a) compared to the expression pattern in dorsal root ganglia innervating other parts of the body (Gerhold et al., 2013). By contrast, the trigeminal ganglia of this species are deficient in the expression of ion channels typically associated with thermosensation (TRPV1, TRPA1, TRPM8). Intriguingly, this same pattern was found in the trigeminal neurons of the highly mechanosensitive bill of tactile-feeding waterfowl (ducks) relative to visually foraging birds (Schneider et al., 2014). It was thus argued that the highly specialised rostra of these species evolved to provide extremely high tactile sensitivity at the cost of reduced thermosensation. However, since temperature sensing ion channels have been implicated in the control of peripheral blood flow in mammals (Fromy et al., 2018), the reduction of temperature sensing ion channels in the trigeminal ganglia, combined with the functional thermoconforming evidence of the star provided herein, suggests an additional explanation. Specifically, the low number of temperature sensing ion channels within the nasal epithelia that have previously been linked to their intense anatomical specialisation for mechanosensation (see Schneider et al., 2016) may also be related to the thermally non-responsive blood supply to the star. In other words, the evolutionary diminution of temperature responding pathways may prevent the reactive thermoregulatory vasomotion of this structure typically observed in peripheral tissues of other endotherms. It should be stressed that, like star-nosed moles, tactile feeding waterfowl that similarly possess low numbers of temperature-sensing neurons in their bill do not exhibit reductions in feeding efficiency in the cold (Schneider et al., 2014). Taken together, these observations suggest that evolutionary reductions in thermosensing ion channels may be a specialization for somatosensory organs that must operate well below core body temperatures. Curiously, an enhanced sensory response has been observed in the rattlesnake pit organ when cold, as it responds more strongly to stimuli at cooler temperatures than at warm temperatures (Bakken et al., 2018; Cadena et al., 2013).

A further explanation for why the star-nose mole shows low vasomotor responses in the star and front limbs might also be related to the already substantial thermal window found in the sparsely haired tail (Fig. 1). The star-nosed mole tail surface temperature response to handling (see Fig. 1D) and changing environmental conditions correspond closely to that seen in rodent tails (Johansen, 1962; Romanovsky et al., 2002), whereby handling and cold-exposure induces vasoconstriction and warm exposure induces vasodilation (Rand et al., 1965; Vianna and Carrive, 2005). However, the tail of star-nosed moles is unusual among talpids in that it is relatively long and accumulates extensive fat stores during the fall and winter (Hamilton, 1931; Petersen and Yates, 1980). Accordingly, this poorly insulated and high-surface-area appendage is well suited to serve as the primary peripheral thermal window involved in adjustable heat exchange. The eye temperature response, on the other hand provides a unique perspective into the species functional morphology. The eyes of star-nosed moles are minute, have tiny optic nerves, and are likely only used for light/dark discrimination (Catania, 1999; Petersen and Yates, 1980). However, eye surface temperatures were elevated and remained nearly constant across all tested temperatures, demonstrating a continuous and high level of blood flow to an organ that has been argued to serve only a minor contribution to overall sensory input.

A final unresolved question pertains to the mechanism underlying the poikilothermic nature of the star. For example, while the reduction of thermosenstive neurons innervating the star may in part underlie the lack of temperature-dependent vasomotion observed in this study, it does not provide insights into how the star is able to achieve relative thermoconformity with environmental temperatures. While it is possible this ability arises via the operation of countercurrent heat exchangers in the rostral region, presence of these structures have not been identified in previous anatomical studies of this species. Alternatively, this trait may result from intermittent blood flow to the star arising from nasal ray movements. For instance, the nasal rays are oriented perpendicular to the nose while foraging (Fig. 1A, B) though these structures are shifted more or less parallel to the nose when the head is raised (Fig. 1K) and while the mole is inactive. When not foraging, star-nosed moles were also routinely observed to exhibit repetitive flexing and extension of the nasal rays and to ‘groom’ the star with the forepaws. While not definitive, the latter behaviour coincided with a sudden increase in blood flow during one of our experiments (Movie 3) and may be important for promoting blood flow to through the large nasal sinuses in the nasal appendages. These competing mechanisms provide potentially fruitful avenues of research on the thermal biology of this unusual insectivore.

## Conclusions

Unconventionally for a peripheral appendage, the nasal rays of the star-nosed mole show little evidence of reactive vasodilation that other mammalian appendages often demonstrate. This thermoconforming response may be related to the high energetic consequences of heat dissipation in a typical peripheral tissue that would accompany the active foraging lifestyle of the star-nosed mole. Since they spend much of their life foraging in environments of high thermal conductance, any body heat reaching the star would be rapidly dissipated to the environment. Extending these results to other “sensory specialists” could be of great interest. For example, numerous ducks have highly sensitive mechanosensation in the bill yet still forage at variable water temperatures (Schneider et al., 2014), the duck-billed platypus (*Ornithorhynchus anatinus*) relies on an extensively innervated and presumably well vascularised bill for electroreception to forage underwater (Scheich et al., 1986), while the echidna (*Tachyglossus aculeatus*) bill is a specialised mechanosensory appendage (Proske et al., 1998). Whether these appendages also demonstrate heat conservation through promiscuous vasoconstriction while foraging is unknown, but the discovery of similar or divergent responses would shed light on the physiological conservatism between how thermoregulatory vascular control has evolved to minimise influences on the sensory systems.

## Acknowledgements

We thank Josh Campbell for assistance with mole capture, and the British Broadcasting Corporation Studios Natural History Unit for accommodating this study. This research was supported by NSERC Discovery Grants to GJT (RGPIN-2020-05089) and KLC (RGPIN-2016-06562) and an NSERC Research Tools and Instrumentation Grant to GJT (NSERC RTI-2021-00278)

## Data availability

http://hdl.handle.net/10464/16980

## Author Contributions

Conceptualization: GJT and KLC; Investigation: GJT and KLC; Formal Analysis: GJT; Writing – Original Draft Preparation: GJT and KLC; Writing – Review & Editing: GJT and KLC; Visualization: GJT; Data Curation: GJT.; Funding Acquisition: GJT and KLC.

## Supplementary Material

**Movie 1.**
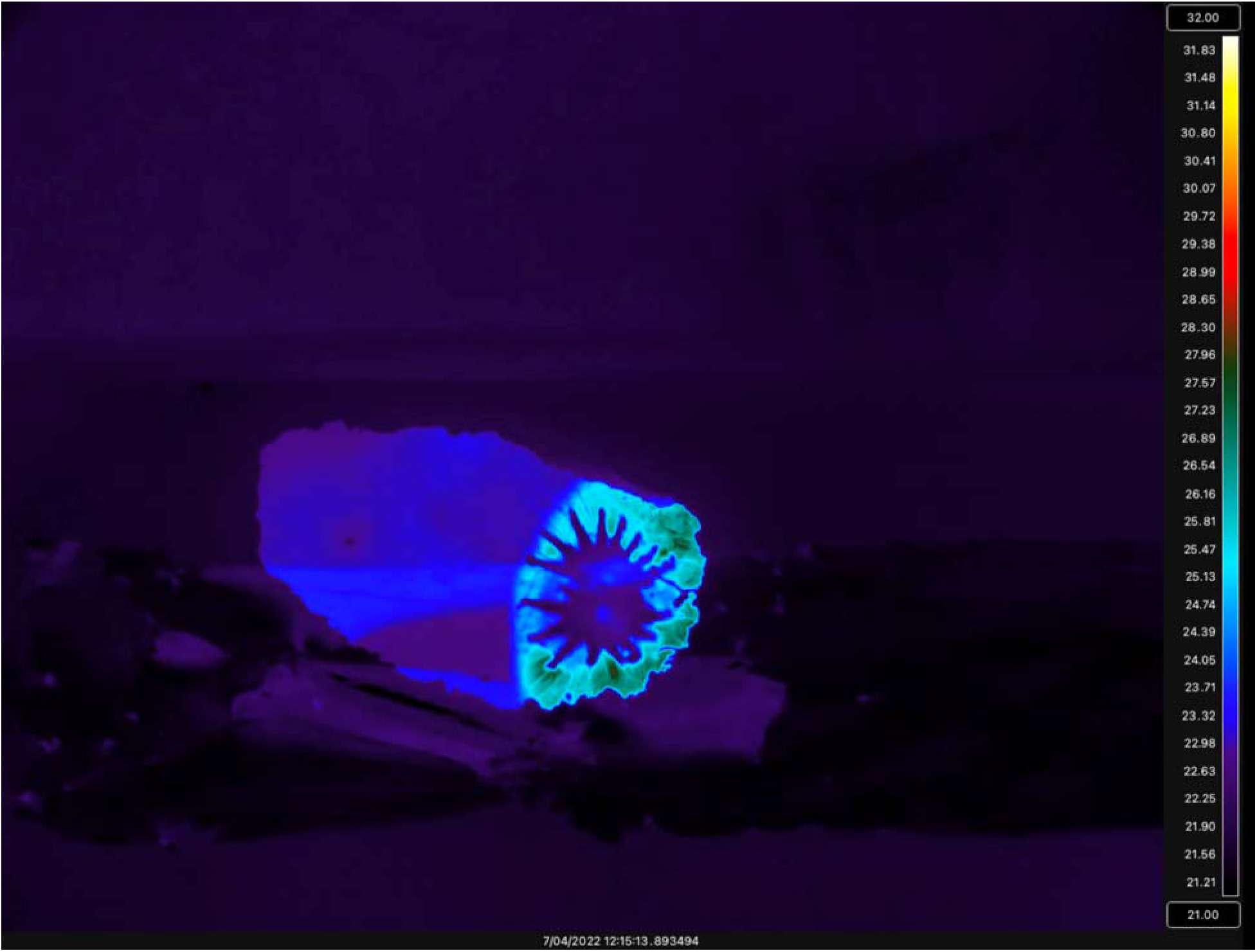
Thermal video (48 frames/sec) of a star-nosed mole scanning an open surface, passing over this surface in 0.48 seconds. The ‘window’ surface was a piece of plastic (Saran™ wrap) stretched over a hole within an artificial tunnel environment.

**Movie 2.**
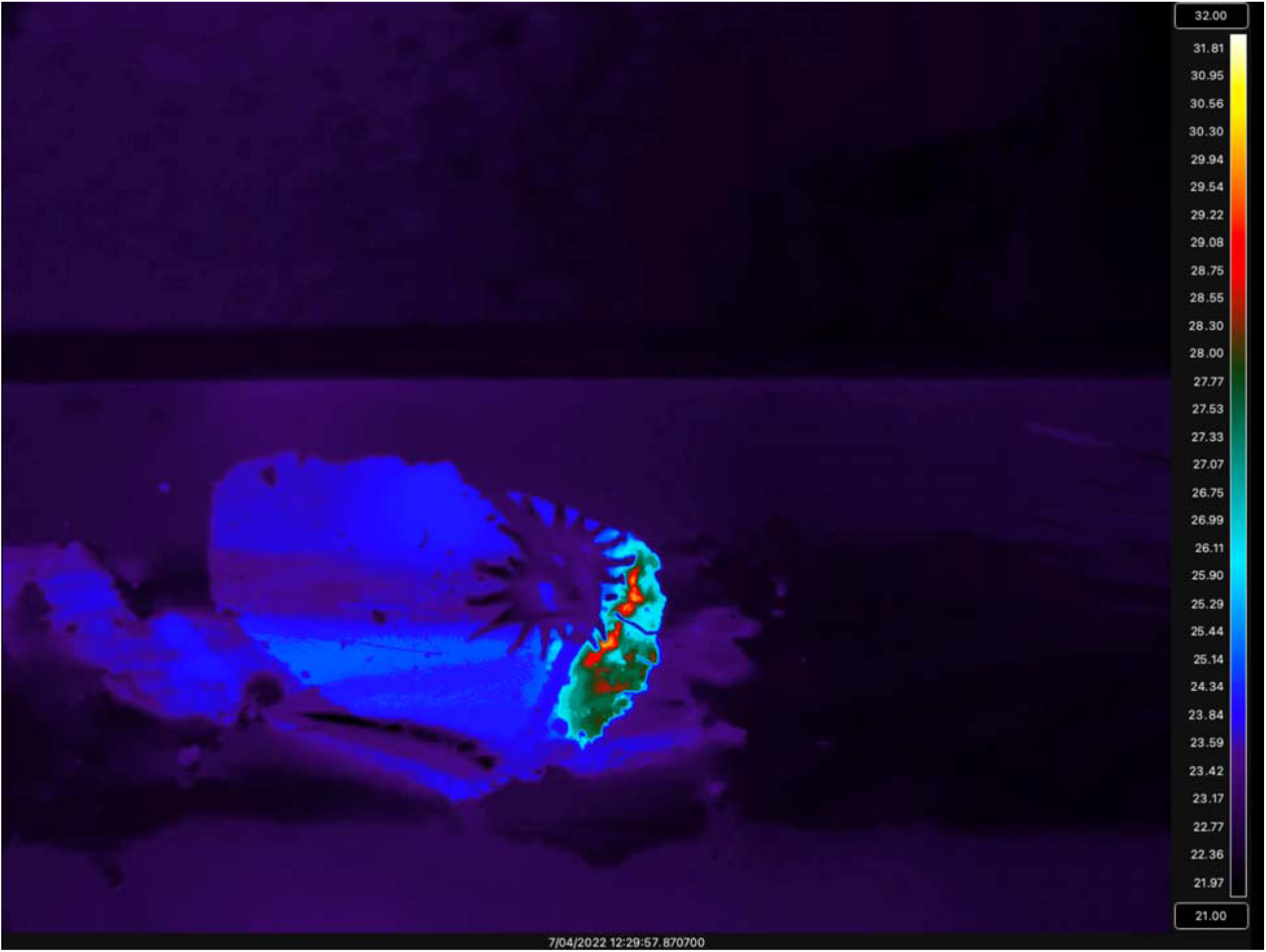
Thermal video (48 frames/sec) of a star-nosed mole scanning an open surface containing a small earthworm, which was detected and consumed in under 3 seconds. The ‘window’ surface was a piece of plastic (Saran™ wrap) stretched over a hole within an artificial tunnel environment.

**Movie 3.**
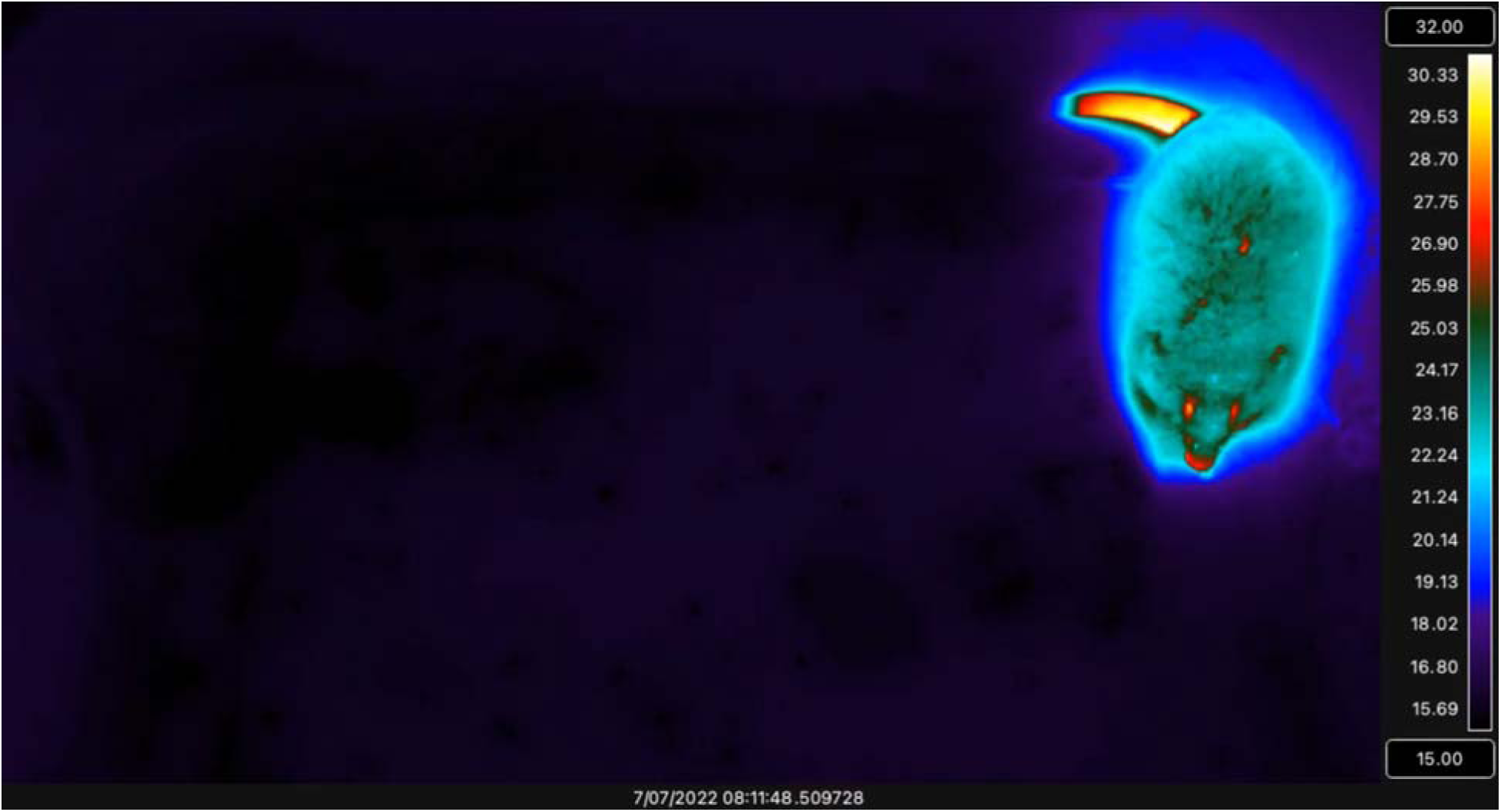
Time-lapse thermal video (frame rate 1 Hz, playback rate 10 Hz) of a star-nosed mole grooming its nose. The star is not splayed open during grooming, and part-way through the video (Timestamp 08:11:48.5, UTC +0), a sudden rise in nasal ray temperature is evident. Since the front limbs are also warm during this video, it is not clear if heat is being transferred from the limbs or if this arises from vasodilation of the nasal rays.

